# The HDAC2-selective inhibitor KTT-1 attenuates autoimmune arthritis by inhibiting osteoclast differentiation in mice

**DOI:** 10.1101/2025.10.20.683312

**Authors:** Yumi Ogihara, Wei Zhe, Yotaro Kodaira, Toshifumi Tojo, Yukihiro Itoh, Takayoshi Suzuki, Daisuke Takahashi, Koji Hase

## Abstract

Histone deacetylases (HDACs) are crucial epigenetic regulators of gene expression through the reversible acetylation of lysine residues, and their inhibition is known to exert anti-arthritic effects. Given the critical role of osteoclasts in bone destruction in rheumatoid arthritis (RA) pathogenesis, targeting HDAC2—whose expression is upregulated during osteoclast differentiation—represents a promising therapeutic strategy. Although the development of highly specific HDAC2 inhibitors has recently attracted considerable attention, no such inhibitor has yet been successfully developed. In the current study, we found that the kinetic-selective HDAC2 inhibitor KTT-1 effectively suppressed arthritis symptoms in the collagen-induced arthritis (CIA) mouse model without causing severe side effects. Furthermore, KTT-1 inhibited osteoclast differentiation at an early stage by downregulating c-Fos expression, suggesting that the KTT-1-mediated anti-arthritis effect was achieved by inhibiting osteoclast differentiation. These findings highlight KTT-1 as a promising therapeutic candidate for the development of targeted synthetic disease-modifying antirheumatic drugs (tsDMARDs).

## Introduction

Rheumatoid arthritis (RA) is a chronic systemic autoimmune disease characterized by synovitis and the progressive destruction of cartilage and bone (Smolen and Steiner, 2003). RA-associated bone loss manifests in several forms, including local erosions in the inflamed joints, periarticular bone loss, and systemic osteoporosis and osteopenia (Zerbini et al., 2017; Steffen et al., 2019). This bone destruction is driven by an imbalance in bone remodeling, characterized by enhanced osteoclast differentiation and activity, which accelerate bone resorption, coupled with a relative suppression of osteoblast-mediated bone formation(Choi et al., 2009; Komatsu and Takayanagi, 2022). Consistent with these findings, mice lacking receptor activator of nuclear factor-κB ligand (RANKL), a crucial factor in osteoclast differentiation, are protected against bone erosion in an experimental model of RA (Pettit et al., 2001). Furthermore, patients with RA exhibit increased numbers and activation of osteoclasts, which significantly contribute to joint damage (Shigeyama et al., 2000). Despite the critical role of osteoclasts in the pathogenesis of rheumatoid arthritis (RA), therapeutic strategies directly targeting osteoclast differentiation to prevent bone destruction remain limited.

The genetic information in eukaryotic cells is packaged into chromatin by wrapping DNA around histone core proteins(Hassig et al., 1998). Histone deacetylases (HDACs) are a family of enzymes that play a central role in the regulation of gene expression by catalyzing the removal of acetyl groups from lysine residues on histone tails and non-histone proteins (West and Johnstone, 2014). This deacetylation typically results in chromatin condensation and the repression of gene transcription (Gallinari et al., 2007). In humans and mice, HDAC isozymes are classified into four main groups: class I (HDAC1, HDAC2, HDAC3, and HDAC8), class IIa (HDAC4, HDAC5, HDAC7, and HDAC9), class IIb (HDAC6 and HDAC10), and class IV (HDAC11) (Subramanian et al., 2010). There are also NAD^+^-dependent class III sirtuins (SIRT1-7). While broad-spectrum pan-HDAC inhibitors (HDACi) have anti-arthritis effects in preclinical mouse models(Chung et al., 2003; Lin et al., 2007), their therapeutic potential is greatly limited by pronounced adverse effects—including fatigue, anorexia, diarrhea, vomiting, weight loss, and alterations in serum biochemical markers—arising from their lack of selectivity and consequent activity against multiple HDAC isozymes beyond the intended target (Subramanian et al., 2010; Faulkner et al., 2019). This has underscored the need to develop isozyme-specific HDACi capable of selectively targeting distinct HDAC functions to suppress osteoclast differentiation and thereby prevent bone destruction in RA, while minimizing systemic toxicity.

Although the HDAC isozymes most effective in inhibiting arthritis are still under investigation, several studies have demonstrated that class I HDACs are upregulated in the synovial fibroblasts and tissues from both rheumatoid arthritis patients and animal models (Kawabata et al., 2010; Meng et al., 2020; Mao et al., 2023). Consistent with these findings, inhibition of class I HDACs, particularly HDAC1 and HDAC2, has been shown to ameliorate arthritis symptoms and attenuate disease progression in arthritis models(Hawtree et al., 2015; Faulkner et al., 2019; Algate et al., 2020). We previously demonstrated that a HDAC1-selective inhibitor suppresses cytokine production in macrophages and alleviates symptoms in collagen-induced arthritis (CIA) and collagen antibody-induced arthritis (CAIA) models in mice (Zhe et al., 2022). However, this inhibitor exhibited only marginal suppression of osteoclast differentiation, even at high concentrations, suggesting that HDAC1 may plays only a minor role in osteoclast differentiation. In contrast, other studies have shown that HDAC2 expression is upregulated during osteoclastogenesis, and that the genetic deletion of HDAC2 in bone marrow-derived macrophages impairs osteoclast differentiation (Dou et al., 2016). These findings indicate that HDAC2 is essential for the differentiation of osteoclasts and represents a promising therapeutic target for the development of novel treatments in RA aimed at preventing bone destruction.

In the current study, we investigated the therapeutic potential of KTT-1, a kinetically selective inhibitor of HDAC2 (Mishima et al., 2023), in experimental models of autoimmune arthritis. KTT-1 markedly attenuated the disease severity in both the CIA and CAIA.

Furthermore, KTT-1 inhibited the differentiation of bone marrow-derived osteoclasts *in vitro*, likely by downregulating c-Maf expression during the early phase of osteoclastogenesis. Collectively, these findings demonstrate that selective HDAC2 inhibition by KTT-1 represents a promising strategy for the development of targeted synthetic disease-modifying antirheumatic drugs (tsDMARDs) aimed at preventing bone destruction in rheumatoid arthritis.

## Materials and methods

### 2.1 Mice

C57BL/6J Jcl mice and DBA/1JJmSlc mice were purchased from CLEA Japan (Tokyo, Japan) and Sankyo Labo Service (Tokyo, Japan), respectively, and maintained under specific pathogen-free conditions. They were fed CE-2 pellet chow (CLEA Japan). All mouse experiments were performed in accordance with protocols approved by the Keio University Institutional Animal Care and Use Committee.

### 2.2 Bone marrow-derived macrophages

Bone marrow (BM) cells were isolated from the femur and tibia of male C57BL/6J Jcl mice (6–8 weeks old). After erythrocyte lysis treatment, the BM cells were cultured in a differentiation medium, namely, high-glucose Dulbecco’s Modified Eagle Medium (DMEM; Nacalai Tesque, Kyoto, Japan) with 10% heat-inactivated fetal bovine serum (FBS) (MP Biomedicals, Irvine, CA), penicillin/streptomycin mixed solution (P/S, 1:100; Nacalai Tesque), β-mercaptoethanol (1:1000, 55 µM; Thermo Fisher Scientific, Waltham, MA), and hydroxyethyl piperazine ethanesulfonic acid buffer solution (HEPES, 10 mM; Nacalai Tesque). Macrophages were polarized toward an inflammatory (M1) phenotype by stimulation with 100 ng/mL lipopolysaccharide (LPS; *E. coli* O111125-05181; Wako, Osaka, Japan), followed by 6h stimulation with 20 ng/mL nigericin (InvivoGen, San Diego, CA). After culturing under polarized conditions, cell supernatants were collected for enzyme-linked immunosorbent assay (ELISA).

### 2.3 Measurement of KTT-1 by LC-MS/MS

KTT-1 in plasma was quantified by a LCMS-8050 tandem mass spectrometry (Shimadzu, Kyoto, Japan) coupled to high-performance liquid chromatography (Shimadzu). To confirm the plasma concentration of KTT-1, mice were administered KTT-1 in their drinking water for 10 days and then euthanized. Plasma samples were deproteinized with an equal volume of acetonitrile containing 100 nM TTA03-107 (Zhe et al., 2022) and centrifuged, and the supernatant was subjected to LC-MS/MS. Chromatographic separation was performed on a Scherzo SS-C18 column (3.0 mm i.d. × 150 mm, 3 μm; Imtakt, Kyoto, Japan) at 40°C with a gradient of mobile phases A: 0.1 % formic acid in water and B: acetonitrile and 100 mM ammonium formate in water (2:3, v/v). The gradient of mobile phase B began at 50% for 3 min at a flow rate of 0.1000 mL/min. Mass spectrometric detection was performed by multiple reaction monitoring (MRM) in the electrospray ionization (ESI) positive ion mode, using m/z of Q1/Q3: 326.05/145.05 and 326.05/117.10.

### 2.4 CIA Model

Six-to seven-week-old male DBA/1JJmsSlc mice were immunized with 50 µL (2 mg/mL) of immunization-grade bovine type II collagen (CII; Chondrex, Woodinville, WA) emulsified in 50 µL (1 mg/mL) of complete Freund’s adjuvant (Chondrex) at the base of the tail on day 1. On day 21, the mice were injected with bovine CII emulsified in incomplete Freund’s adjuvant (Chondrex) as a booster. Then, mice were randomly divided into three groups of 12 mice each: control DMSO, VPA, and KTT-1 groups. KTT-1 and sodium valproate (VPA; FUJIFILM Wako Pure Chemical Corporation, Osaka, Japan) were initially dissolved in dimethyl sulfoxide (DMSO) to prepare stock solutions. These stock solutions were subsequently diluted in MediDrip Sucralose (Clear H2O; Westbrook, ME) to a final concentration of 0.03 mg/mL. Vehicle DMSO was diluted in MediDrip Sucralose at 0.003%. These were orally administered as drinking water starting on the day of booster immunization. All mice were dissected on day 43.

### 2.5 CAIA Model

To prepare the CAIA model, we initially purified anti-collagen Immunoglobulin G (IgG) from the serum of CIA model mice, using PROTEUS Protein G Midi Kit (Protein Ark, Sheffield, US) and Amicon Ultra-15 Centrifugal Filter Units (100 kDa, Millipore, Burlington, MA) following the manufacturer’s instructions. Briefly, serum was collected from CIA mice and filtered through a 0.2 mm syringe filter. It was then diluted 1:1 with Binding Buffer G. Next, 5 mL of Binding Buffer G was added to a spin column and centrifuged at 500 × *g* for 3 minutes to equilibrate the column. The diluted serum was loaded onto the spin column and centrifuged at 150 × *g* for 30 minutes. After washing the column, it was placed into a new tube containing Neutralization Buffer. Elution Buffer was added to the column and centrifuged at 500 × *g* for 3 minutes to elute the IgG. This elution step was repeated once. After pre-rinsing the filter with MilliQ water, serum was added up to a volume of 15 mL and centrifuged at 5,000 × *g* for 30 min. After centrifugation, the concentrated sample was collected, and the protein concentration was measured and adjusted to 50 mg/mL. The purified samples were stored at −80°C until use. Six-to seven-week-old male DBA/1JJmsSlc mice were randomly divided into four groups, each consisting of 10 mice, on day 0: control DMSO, VPA, and KTT-1 groups. To induce CAIA, 5 mg of IgG purified from CIA mice was administered intraperitoneally on day 0, followed by an intraperitoneal injection of 50 µg of LPS (*E. coli* O111; FUJIFILM Wako Pure Chemical) on day 3. KTT-1 and VPA were suspended in MediDrip Sucralose and administered orally as drinking water at a concentration of 0.03 or 8 mg/mL, respectively, starting on day 1. All mice were dissected on day 22.

### 2.6 Clinical Evaluation of Arthritis

Clinical parameters were measured using an articular index score. Each paw was scored according to an arthritis scoring system (0, normal; 1, red and swollen toe joint; 2, swollen toe joint and ankle; 3, swelling of the paw; and 4, swelling of all the feet, including the ankle joint. The sum of the scores of the four paws was expressed as the clinical score, with a maximum of 16 points.

### 2.7 Histological Analysis

For histological analysis, after removing the skin, the skinless paws were fixed in 10N Mildform (FUJIFILM Wako Pure Chemical), decalcified in ethylenediaminetetraacetic acid buffer for 15 days, and then embedded in paraffin. The tissue cross-sections were stained with hematoxylin and eosin (H & E; SKK Organization Science Research Institute, Yokohama, Japan). The histological changes were examined under a microscope by blind observers. Synovial inflammation, bone erosion, cartilage damage, and leukocyte infiltration were assessed using a three-parameter scoring system, in which individual scores were summed: joint inflammatory cell infiltration, synovial hyperplasia, and cartilage and bone destruction (0: normal, 1: mild, 2: moderate, 3: marked, and 4: severe). The histological score was determined by summing these scores (Oh et al., 2017).

### 2.8 ELISA

BM cells were cultured as described in section 2.2. Briefly, cells were maintained for 3 days, followed by the addition of KTT-1 and further incubation for 2 days. Subsequently, the cells were stimulated for 10 h either with or without 100 ng/mL LPS to provide an environment for differentiation into inflammatory macrophages (M1) or to maintain an unpolarized state (M0), respectively. The cell supernatants were collected to measure the levels of IL-1β, TNF-α, and IL-17A using ELISA kits (all from BioLegend, San Diego, CA) in accordance with the manufacturer’s instructions.

### 2.9 Osteoclast differentiation assay

BM cells were isolated from the femur and tibia of male C57BL/6J Jcl mice (6–8 weeks old; Clea Japan). Erythrocytes were lysed, and the surviving cells were cultured in a differentiation medium: α-MEM (Gibco) with 10% heat-inactivated fetal bovine serum (FBS) (MP Biomedicals, Irvine, CA), P/S (100×; Nacalai Tesque), β-mercaptoethanol (1:1000, 55 µM; Thermo Fisher Scientific), and HEPES (10 mM; Nacalai Tesque). To induce the osteoclast differentiation, the cells were cultured in complete α-MEM containing 20 ng/Ml recombinant mouse macrophage colony-stimulating factor (rmM-CSF; BioLegend) for 3 days. To determine the effects of KTT-1, the cells were seeded into 96-well plates and cultured with complete α-MEM containing 20 ng/mL M-CSF and 50 ng/mL RANKL (BioLegend) with or without KTT-1 at 37 °C in a 5% CO2 incubator. The culture medium was replaced every 2 days. On day 11 of culture, cells were fixed and stained for tartrate-resistant acid phosphatase (TRAP) activity using a TRAP staining kit (Cosmo Bio, Tokyo, Japan), in accordance with the manufacturer’s protocol. TRAP-positive multinucleated cells containing three or more nuclei were counted as osteoclasts.

### 2.10 RNA-sequencing

Bulk 3’ mRNA sequencing was performed on cells at day 6 (before RANKL treatment) and at days 8 and 10 in the osteoclast culture system for KTT-1-treated and control groups. Total RNA was extracted from the cultured osteoclasts using the NucleoSpin RNA Plus kit (740984.250, MACHEREY-NAGEL, Düren, Germany). A cDNA library was synthesized using Collibri 3’ mRNA Library Prep Kit for Illumina Systems (A38110024, Thermo Fisher Scientific) according to the manufacturer’s protocol. The libraries were subsequently purified and enriched with AMPure XP beads (Beckman Coulter, Brea, CA) and then sequenced on an Illumina NovaSeq X Plus with 300 bp paired end reads. FASTQ files were mapped with STAR (version 5.0.1) (Dobin et al., 2012) to the mouse mm10 reference genome with default parameters. Differential gene expression analyses were performed in R using RStudio (RStudio Inc.). For Principal components analysis (PCA), calculation of inter-sample Pearson correlation Raw, non-normalized counts were imported into R and analyzed using the DESeq2 package (http://www.bioconductor.org/packages/release/bioc/html/DESeq2.html) (Love et al., 2014) . Genes with fewer than 10 counts across all samples were excluded, and normalization was performed using default parameters for estimating size factors and dispersions. Genes with a q value < 0.05 and Log2 Fold Change > 1 were defined as “differentially expressed genes” and carried forward for further analysis. Volcano plots were generated with the R package EnhancedVolcano(https://github.com/kevinblighe/EnhancedVolcano) (Blighe et al., 2025). Count per million (CPM) were calculated from the obtained DEG results using edgeR (https://bioconductor.org/packages/release/bioc/html/edgeR.html) (Chen et al., 2025), and biomaRt (https://bioconductor.org/packages/release/bioc/html/biomaRt.html) (Durinck et al., 2005, 2009). Gene Ontology (GO) Biological Process was performed using the Database for Annotation, Visualization, and Integrated Discovery (DAVID; https://david.ncifcrf.gov/) (Huang et al., 2009). GO categories in the biological process ontology were identified based on a significance threshold of q-value < 0.05 and |Log2 (Fold Change) | > 1.

### 2.11 Statistical analysis

Statistical analyses were performed using GraphPad Prism 10 software (GraphPad Software, La Jolla, CA, USA). A *p*-value < 0.05 was considered statistically significant. The specific statistical test used for each experiment is detailed in the corresponding figure legend. In general, for comparisons between two groups, a two-tailed Student’s *t*-test was used for data assumed to have equal variances, while Welch’s t-test was applied for data with unequal variances. For non-normally distributed data, the non-parametric Mann-Whitney U test was used. For comparisons among three or more groups, one-way or two-way analysis of variance (ANOVA) was performed. Post-hoc multiple comparisons were conducted using Tukey’s test (for all pairwise comparisons), or Dunnett’s test (for comparisons against a single control group), as appropriate. Data are typically presented as mean ± s.d. The experiments were not randomized, and the investigators were not blinded to allocation during experiments or outcome assessment.

## 3. Results

### 3.1. Oral Administration of KTT-1 Attenuates Arthritis Severity in CIA Model

To measure the systemic exposure of KTT-1 following oral administration, we first conducted a pharmacokinetic assessment. DBA/1J mice were administered the KTT-1 in drinking water for 10 consecutive days. Analysis of plasma samples collected thereafter showed a median (interquartile range) KTT-1 concentration of 356.0 (320.5–576.1) nM, confirming successful systemic delivery and sustained exposure via oral administration (Fig. 1A). Subsequently, we investigated the therapeutic efficacy of KTT-1 in the CIA model. Following the booster immunization with CII, DBA/1J mice were treated with KTT-1 in drinking water (approximately 6 mg/kg/day) or VPA (approximately 1,200 mg/kg/day), a non-selective HDAC inhibitor known to target multiple Class I and Class II HDACs, as a reference for broad HDAC inhibition (Göttlicher et al., 2001). A control group received regular drinking water. Throughout the treatment period, general health was monitored by assessing body weight. Mice treated with KTT-1 exhibited a rate of body weight gain comparable to or slightly better than that of the control group, whereas the VPA-treated group showed a tendency towards reduced weight gain, suggesting better tolerability of KTT-1 at the tested dose (Fig. 1B). Importantly, evaluation of clinical arthritis scores demonstrated that KTT-1 treatment significantly attenuated the severity of arthritis compared with the control group, whereas VPA only slightly improved clinical score (Fig. 1C). Histopathological examination of joint tissues further supported the protective effect of KTT-1. Joints from control CIA mice displayed typical pathological features of severe arthritis, including synovial hyperplasia, extensive inflammatory cell infiltration into the synovial tissue and joint space, pannus formation, and significant cartilage erosion and bone destruction. In contrast, joints from KTT-1-treated mice showed markedly reduced synovial inflammation and immune cell infiltration, with preservation of normal joint architecture (Fig. 1D, E).

**Fig. 1.**
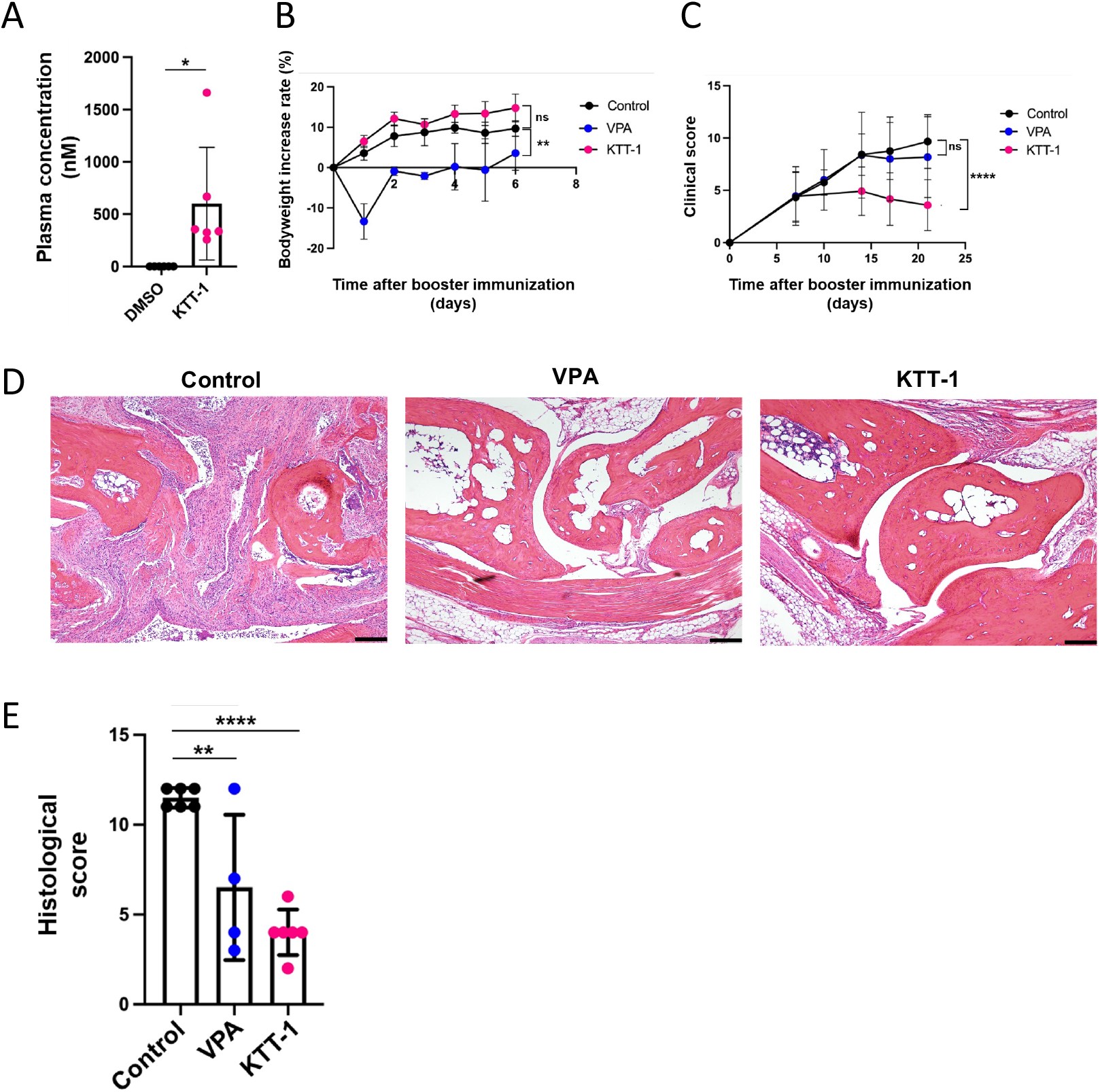
KTT-1 alleviated the severity of arthritis in CIA model mice Plasma concentration of KTT-1 in mice treated with KTT-1 for one week (n=6/group, unpaired *t*-test) (A). Body weight increase rates (B) and (C) clinical scores indicate arthritis severity and potential adverse effects by respective treatment in CIA mice (n = 12 mice/group, one-way ANOVA or two-way ANOVA followed by Dunnett’s test, compared to the untreated control group). Representative H&E staining graph of joint sections shows the degree of synovial inflammation, bone erosion, cartilage destruction, and leukocyte infiltration. Bars: 200 µm (D). Histological scores were calculated by H&E staining graph of respective group (n=12 mice/group, Dunnett’s test, compared to the untreated control group) (E). *p<0.05, **p<0.01, ***p<0.001. Data for the control and VPA groups were reproduced from our previously published work (Zhe et al., 2022).

### 3.2 KTT-1 Suppresses Arthritis Severity in the CAIA Model

To investigate whether the therapeutic effect of KTT-1 in the CIA model involved modulation of the underlying autoimmune response, we measured the serum levels of CII-reactive IgG, a driver of the CIA pathogenesis. Administration of either VPA or KTT-1 did not significantly reduce the levels of anti-CII total IgG, IgG1, IgG2a, and IgG2b compared with the control group (Fig. 2A). These data suggest that the therapeutic mechanism of KTT-1 likely acts downstream or independently of the autoantibody production phase in the CIA model. To further investigate whether KTT-1 modulates in the later phase of RA pathogenesis, particularly inflammation driven by autoantibodies and innate immune cells, we employed the CAIA model. In the CAIA model, arthritis is rapidly induced by administering an anti-CII antibody cocktail followed by lipopolysaccharide (LPS), thereby bypassing the need for IgG-inducing adaptive immune responses. DBA/1J mice were administered with KTT-1 following the injection of anti-CII IgG antibodies. Consistent with observations in the CIA model, KTT treatment exhibited normal body weight gain relative to control vehicle treatment, whereas VPA treatment group led to a decrease in body weight (Fig. 2B). KTT-1 treatment significantly alleviated the severity of arthritis in the CAIA model (Fig. 2C). Based on these observations, the therapeutic efficacy of KTT-1 primarily derives from suppression of the downstream inflammatory cascade, rather than attenuation of autoantibody production.

**Fig. 2.**
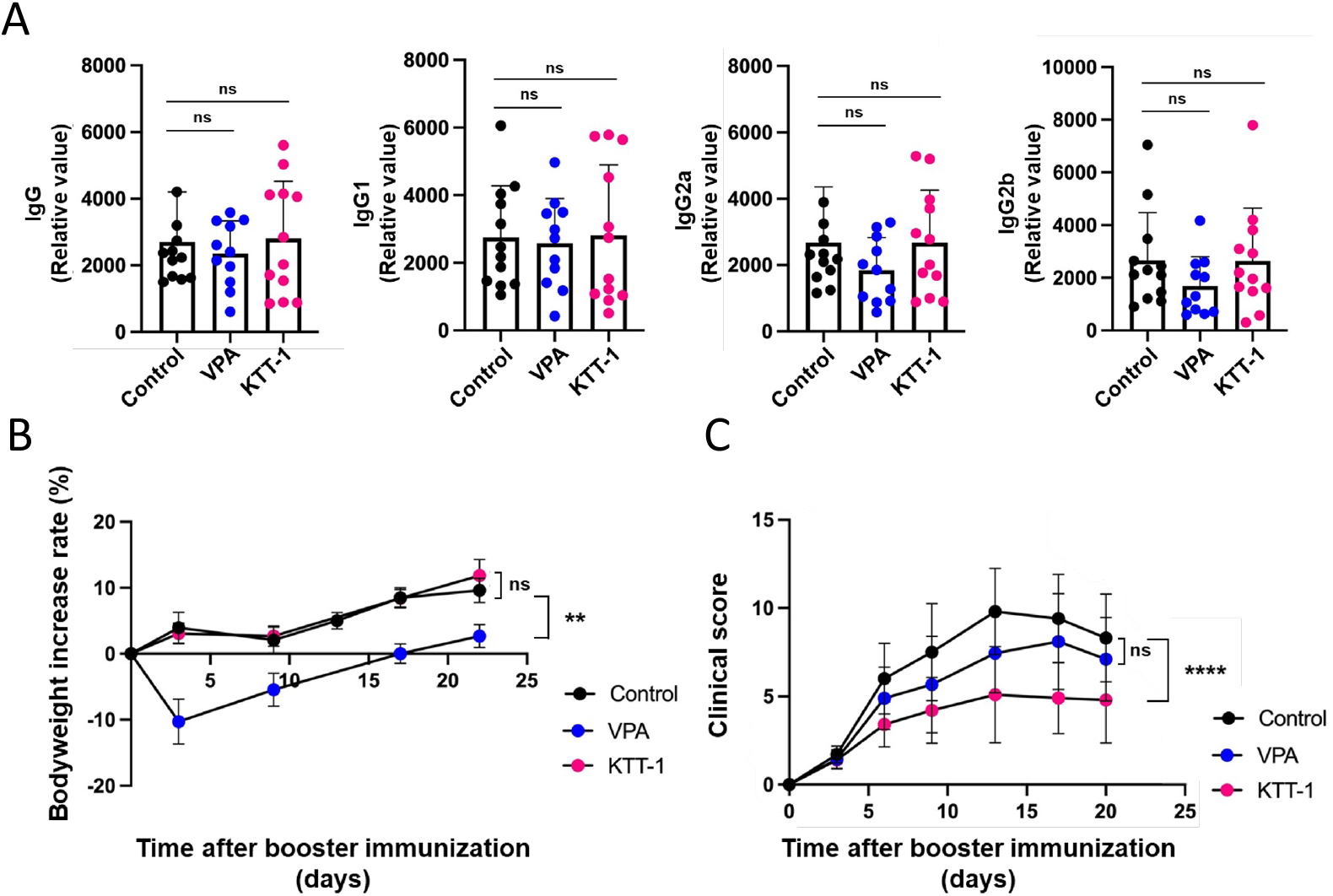
KTT-1 alleviated the severity of arthritis in the CAIA model mice Serum levels of anti-CII IgG, anti-CII IgG1, anti-CII IgG2a, and anti-CII IgG2b in CIA mice (n=12 mice/group, Dunnett’s test, compared to the untreated control group) (E). Body weight increase rate (B) and Clinical score (C) in CAIA mice (n=10 mice/group, Dunnet’s test, compared to the untreated control group). *p<0.05, **p<0.01, ***p<0.001. Data for the control and VPA groups were reproduced from our previously published work (Zhe et al., 2022) .

### 3.3 KTT-1 Primarily Suppresses Osteoclast Differentiation Rather Than Macrophage Cytokine Production *In Vitro*

The pathogenesis of CIA and CAIA involves the production of inflammatory cytokines, largely by macrophages, as well as bone destruction mediated by osteoclasts (Zhe et al., 2022). To dissect the cellular mechanisms underlying the therapeutic effect of KTT-1 in CIA and CAIA models, we investigated its direct actions on these two key cell types *in vitro*. First, we assessed the impact of KTT-1 on pro-inflammatory cytokine production by bone marrow-derived macrophages (BMDMs) upon LPS stimulation. KTT-1 treatment led to a modest reduction in IL-1β production in a dose-dependent manner, whereas significant suppression of IL-6 production was observed only at higher concentrations (Fig. 3). KTT-1 did not significantly affect TNF-α production (Fig. 3). These data suggest that KTT-1 may have a relatively limited capacity to suppress pro-inflammatory cytokine production by macrophages at the concentrations likely achieved *in vivo* (Fig. 1A). Subsequently, we examined the effect of KTT-1 on osteoclast differentiation. BM cells were cultured under BMDM differentiation conditions, followed by osteoclastogenic conditions in the presence of varying concentrations of KTT-1. We observed that KTT-1 markedly inhibited mature, multinucleated osteoclasts in a dose-dependent fashion, as evidenced by TRAP staining and quantification (Fig. 4A, B). Taken together, our findings suggest that KTT-1 exerts its therapeutic effect primarily by inhibiting osteoclast differentiation, with only modest effects on macrophage cytokine production (particularly IL-1β).

**Fig. 3.**
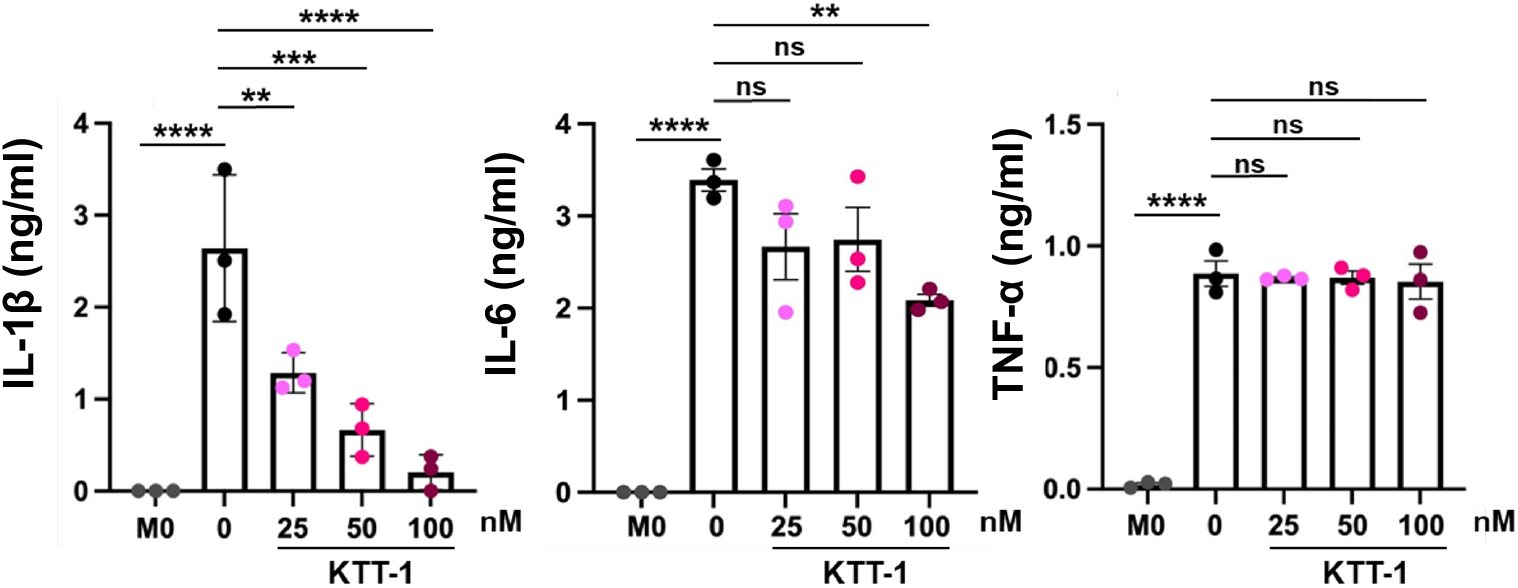
The effects of KTT-1 treatment on the production of inflammatory cytokines by macrophages Effect of KTT-1 on IL-1β, IL-6, and TNF-α production under M0 culture conditions (without LPS stimulation) or M1 culture conditions (with LPS, control group) (n=3, Dunnett’s test, compared to the untreated control group). *p<0.05, **p<0.01, ***p<0.001. Data for the M0 and control groups were reproduced from our previously published work (Zhe et al., 2022) .

**Fig. 4.**
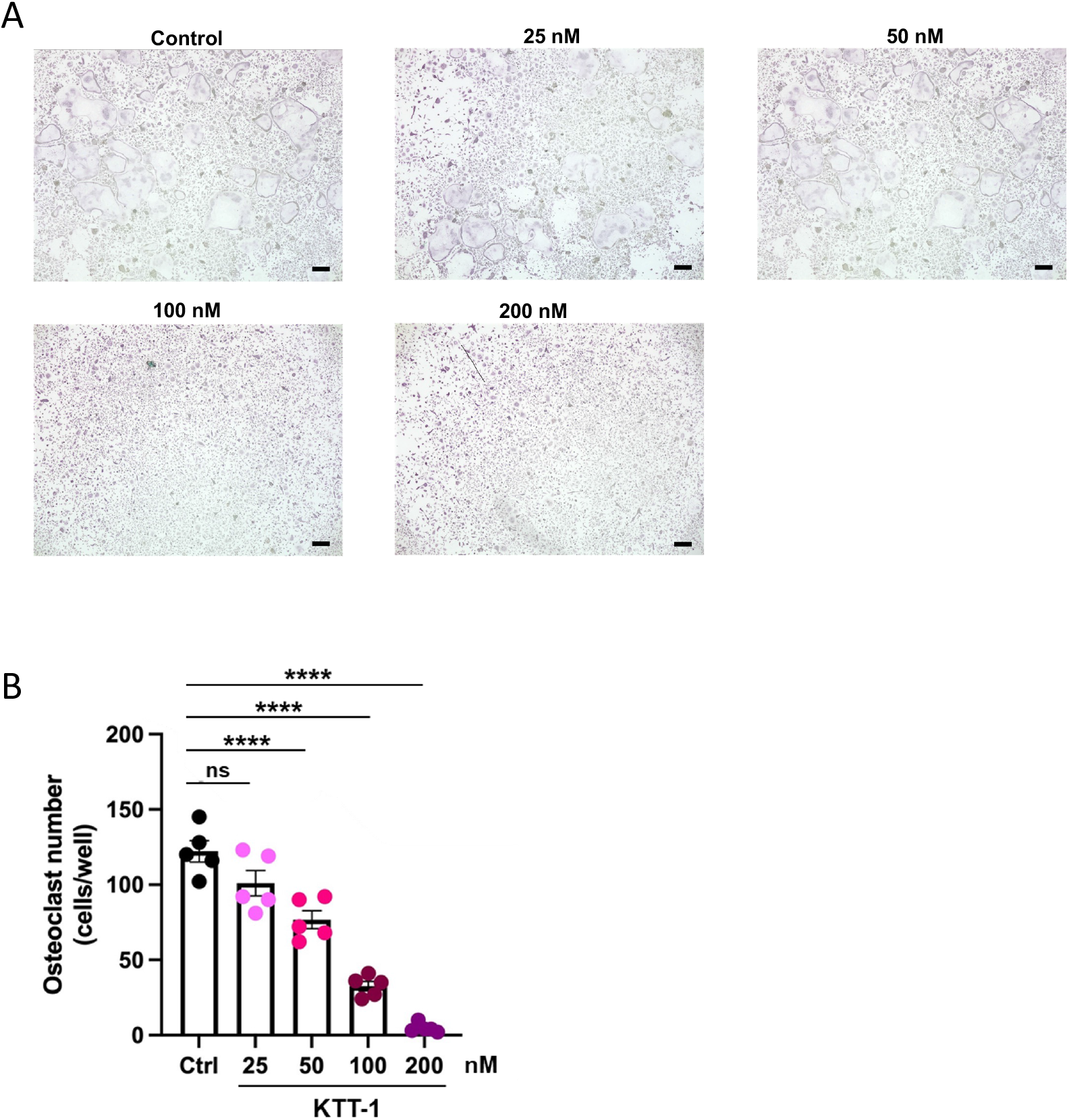
KTT-1 treatment suppressed osteoclast differentiation Induce osteoclasts from in vitro bone marrow cell cultures stimulated with M-CSF and RANKL. Representative microphotograph (A) and quantification of multinucleated TRAP-positive osteoclasts (B) in groups with different concentrations of KTT-1 or without the KTT-1 group. Bars: 200 µm. (n=5/group, Dunnett’s test, compared to the untreated control group). *p<0.05, **p<0.01, ***p<0.001. Data for the control groups were reproduced from our previously published work (Zhe et al., 2022) .

### 3.4 KTT-1 Disrupts Osteoclast Differentiation Progression by Altering Early Stage-Specific Gene Expression

To determine the specific stages of osteoclast differentiation inhibited by KTT-1 and gain insight into the outcomes of HDAC2 inhibition during this process, we performed transcriptome analysis. RNA samples were harvested from the BM-derived macrophages/osteoclasts with or without KTT-1 treatment on day 6 (macrophage), day 8 (the early stage of osteoclast differentiation), and day 10 (the middle to late stages of osteoclast differentiation) (Fig. 5A). PCA of the transcriptome analysis data demonstrated apparent segregation between the KTT-1-treated and control groups on day 10, but not on day 8 (Fig. 5B), indicating a significant impact of KTT-1 on the global gene expression profile during the later stages of osteoclastogenesis. Even though there is no obvious segregation between the groups on day 8, volcano plot analysis identified a substantial number of genes whose expression was significantly altered by KTT-1 treatment (Fig. 5C). Furthermore, gene ontology (GO) enrichment analysis of the gene subset significantly downregulated by KTT-1 treatment on day 8 highlighted biological processes such as “extracellular matrix organization”, “extracellular structure organization”, and “ossification” (Fig. 5D). This suggests that KTT-1 interferes with the expression programs required for osteoclast maturation and function. Indeed, on day 10 of culture, the expression of *Nfatc1*, which serves as the master transcriptional regulator of osteoclastogenesis, as well as *Fos*, which encodes c-Fos, an upstream regulator of Nfatc1, was significantly decreased in the KTT-1 treatment group compared to the control group (Fig. 5E, F).

**Fig. 5.**
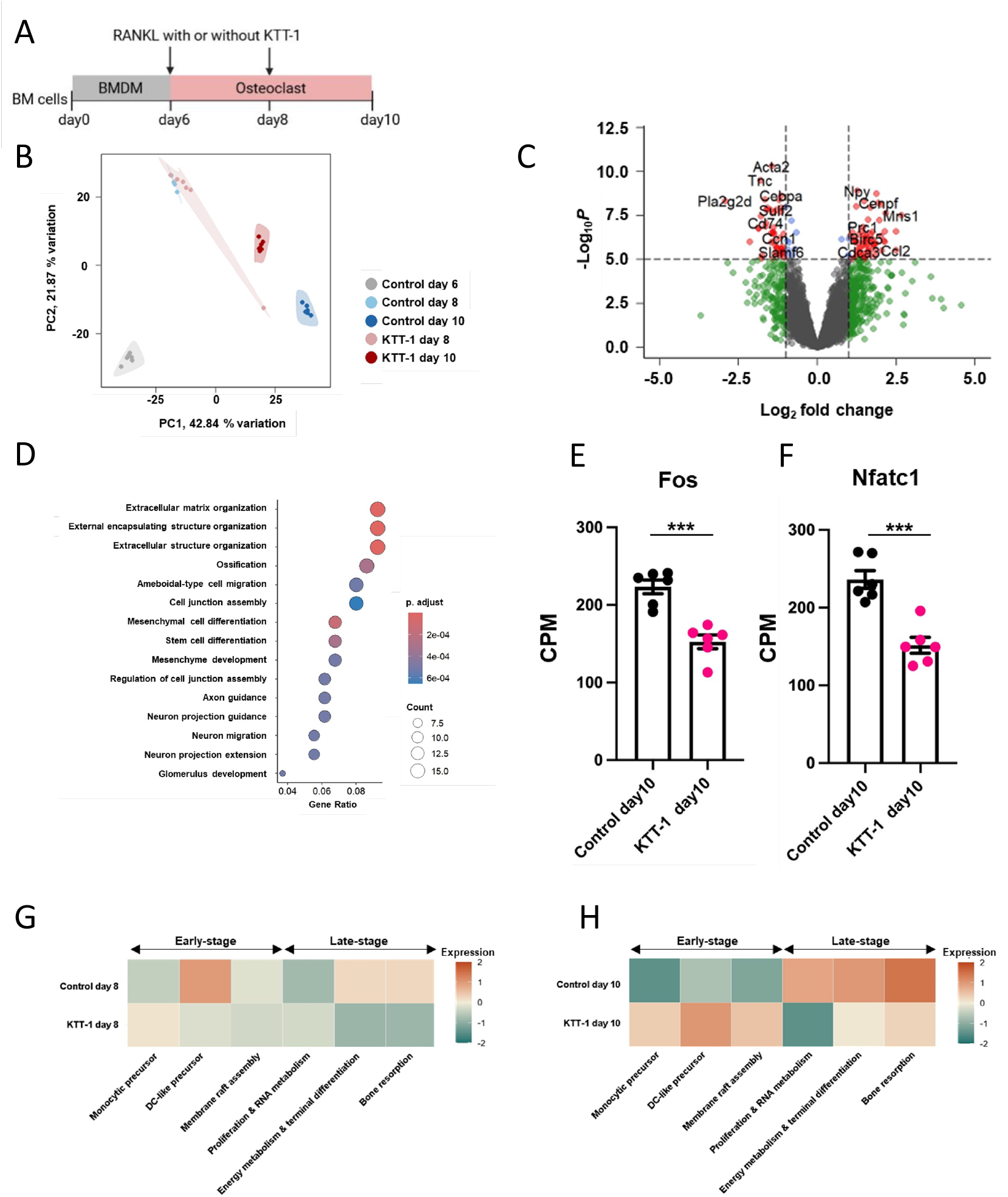
The change of RNA expression by KTT-1 treatment Experimental setup for (B) to (G). Bone marrow (BM) cells were isolated from mice and cultured. RANKL was added on days 6 and 8 of culture to induce differentiation into osteoclasts. In the KTT-1-treated group, KTT-1 was also added on days 6 and 8 of culture (A). Comparison of RNA expression between KTT-1 treated and Control groups by Principal components analysis (B) (n=6/group). Volcano plots comparing KTT-1-treated and control groups at day 8 in vitro culture system (C) (n=6/group). Gene ontology (GO) Biological Process of downregulated genes in KTT-1 treated groups at day 8 (D) (n=6/group). Normalized gene expressed levels (CPM) of *Fos* (E), *Nfatc1* (F), key regulators of osteoclast differentiation. (n=6/group, unpaired *t*-test). *p<0.05, **p<0.01, ***p<0.001. The change of maker gene expression at each step of osteoclast differentiation in KTT-1 treated groups and control groups at day 8 (G) and day 10 (H) (n=6/group).

To more precisely determine the differentiation stage at which KTT-1 exerts its inhibitory effect during the multi-step differentiation process, we examined the expression levels of established marker genes associated with distinct stages of osteoclast differentiation (Tsukasaki et al., 2020). On day 8 of culture, in the control group, marker genes of DC-like precursor were upregulated, indicating progression of early differentiation, whereas KTT-1-treated cells appeared to be arrested at the monocytic precursor stage (Fig. 5G). This pattern was further accentuated by day 10, where KTT-1-treated cells exhibited persistently elevated expression of early-stage markers alongside significantly reduced expression of marker genes of late-stage maturation, such as “proliferation and RNA metabolism”, “energy metabolism and terminal differentiation”, and “bone resorption” (Fig. 5H). These dynamic changes in stage-specific marker gene expression profiles strongly suggest that KTT-1 arrests osteoclast differentiation after the early stages, thereby preventing progression toward terminal differentiation and maturation.

## Discussions

This study explored the therapeutic potential and mechanisms of KTT-1, a recently developed HDAC2-selective inhibitor (Mishima et al., 2023), in preclinical mouse models of RA. Our findings demonstrate that oral administration of KTT-1 significantly ameliorates disease severity in both the T and B-cell-dependent CIA model and the antibody-mediated CAIA model. Notably, this therapeutic efficacy was achieved without the weight loss observed with the non-selective pan-HDAC inhibitor VPA, suggesting a superior therapeutic window and highlighting the potential advantage of isoform-selective HDAC inhibition for RA treatment.

A key aspect of RA pathogenesis involves the interplay between immune-driven inflammation and joint destruction (Firestein, 2003). Our results provide insights into how KTT-1 might interfere with these processes. The lack of a significant impact on systemic anti-CII autoantibody levels in the CIA model, together with its effectiveness in the CAIA model, suggests that KTT-1 mainly targets downstream effector pathways rather than the initial autoimmune activation, including autoantibody production. This supports the idea that KTT-1 exerts its therapeutic effects by reducing the outcomes of established autoimmune responses rather than preventing their initiation.

We compared the effects of KTT-1 on macrophages and osteoclasts in vitro to identify cellular targets. While KTT-1 showed only limited ability to suppress TNF-α and IL-6 production by BMDMs, it strongly inhibited osteoclast differentiation from BM precursors. This is in sharp contrast to our previous findings with an HDAC1-selective inhibitor, TTA03-107, which effectively suppressed cytokine production in BMDMs and subsequent Th17 responses, yet had minimal impacts on osteoclastogenesis (Zhe et al., 2022). This divergence highlights the distinct, non-redundant roles of HDAC1 and HDAC2 in cell types relevant to RA. HDAC1 seems to mainly regulate pathways that induce inflammatory cytokines in macrophages, while HDAC2 appears essential for osteoclast differentiation and maturation. Therefore, selective inhibitors targeting HDAC1 or HDAC2 could provide tailored therapeutic options depending on disease stage: HDAC1 inhibitors may mainly reduce synovial inflammation, whereas HDAC2 inhibitors like KTT-1 might be particularly effective in preventing osteoclast-mediated bone erosion in destructive arthritis.

Our transcriptomic analysis provided further mechanistic insights into the impact of KTT-1 on osteoclastogenesis. KTT-1 treatment significantly alters the gene expression landscape during osteoclast differentiation, leading to an accumulation of early-stage markers and a reduction in late-stage maturation markers. This indicates that KTT-1 likely arrests osteoclast development post-commitment but prior to full maturation, preventing the formation of functional bone-resorbing cells. Consistent with this, GO analysis highlighted downregulation of pathways related to extracellular matrix organization and ossification. As a molecular basis for this differentiation arrest, we found that the expression of NFATc1, the master transcription factor for osteoclastogenesis, was markedly suppressed by KTT-1 treatment. Furthermore, the expression of c-Fos, which plays an essential role in inducing Nfatc1 transcription downstream of RANKL signaling, was similarly suppressed (Asagiri and Takayanagi, 2007). It is known that c-Fos and NFATc1 work cooperatively in response to RANKL signaling to activate numerous osteoclast-specific genes, such as TRAP, cathepsin K, and the calcitonin receptor (Takayanagi et al., 2002; Matsumoto et al., 2004; Kim et al., 2005; Kim and Kim, 2014). Therefore, the concurrent suppression of these two critical transcription factors, c-Fos and Nfatc1, is considered a plausible molecular mechanism for the potent inhibition of osteoclast differentiation by KTT-1. Given that HDAC2 inhibition leads to histone hyperacetylation, it is likely that deacetylation mediated by HDAC2 is essential, either directly or indirectly, for the proper transcriptional activation of key regulators like c-Fos and NFATc1 during osteoclast development. By inhibiting HDAC2, KTT-1 may disrupt this critical epigenetic regulation. However, further experimental validation is required to substantiate this hypothesis.

While our data strongly support osteoclast inhibition as a key mechanism of KTT-1 action, contributions from modulating other cell types (such as synovial fibroblasts, chondrocytes, or specific immune cells) cannot be dismissed. Moreover, although we observed decreased Fos expression, the upstream HDAC2-dependent regulatory elements controlling its transcription are still to be identified. Future epigenetic studies are needed to find direct target genes of HDAC2. Additionally, despite the favorable short-term tolerability of KTT-1 compared to VPA, the long-term safety and effects on systemic bone homeostasis must be carefully evaluated. Lastly, applying these findings from mouse models to human RA will require caution, although the fundamental roles of osteoclasts and HDACs are typically conserved across species.

In conclusion, this study identifies the HDAC2-selective inhibitor KTT-1 as a promising candidate for tsDMARDs. Its distinct mechanism of action, compared with that of the HDAC1 inhibitor, highlights the potential for isoform-selective HDAC inhibitors to target specific pathological processes in RA. KTT-1 may provide a strategy to effectively prevent joint destruction while minimizing the safety concerns linked to pan-HDAC inhibitors. Further research into its detailed molecular mechanisms and long-term effects is warranted.

## Ethics Statement

All animal experiments were approved by the Animal Studies Committees of Keio University.

## Author Contribution

YI, TK, and TS synthesized KTT-1. WZ primarily performed *in vivo* and *in vitro* experiments with the support of YO, DT, and YK. DT and KH designed the study. YO and DT wrote the manuscript. KH revised the manuscript. DT and KH supervised the project.

## Acknowledgments

We thank Tomohiro Nishimura for instructing the LC-MS/MS analysis and Koichiro Suzuki and Shunsuke Kimura for valuable discussions. We are also grateful to the animal facility at the Keio University Faculty of Pharmacy for breeding and maintaining mouse strains.

## Scope Statement

We demonstrated that the HDAC2-selective inhibitor KTT-1 effectively suppresses arthritis symptoms by inhibiting osteoclast differentiation. Although HDAC inhibitors have anti-arthritis effects, the clinical application of non-specific inhibitors is limited due to side effects. Our findings suggest that KTT-1 could be a novel therapeutic agent for rheumatoid arthritis.

